# The Hippo pathway effector YAP inhibits NF-κB signaling and ccRCC growth by opposing ZHX2

**DOI:** 10.1101/2024.06.21.600079

**Authors:** Xu Li, Yong Suk Cho, Yuhong Han, Mengmeng Zhou, Yuchen Liu, Yingzi Yang, Shu Zhuo, Jin Jiang

## Abstract

The prevailing view in the cancer field is that Hippo signaling pathway functions as a tumor suppressor pathway by blocking the oncogenic potential of the pathway effectors Yes1 associated transcriptional regulator (YAP)/transcriptional coactivator with PDZ-binding motif (TAZ). However, YAP can also function as a context-dependent tumor suppressor in several types of cancer including clear cell renal cell carcinomas (ccRCC). We find that, in additional to inhibiting hypoxia-inducible factor 2α (HIF2α), a major oncogenic driver in *Von Hippel-Lindau* (*VHL*)*-/-* ccRCC, YAP also blocks nuclear factor κB (NF-κB) signaling in ccRCC to inhibit cancer cell growth under conditions where HIF2α is dispensable. Mechanistically, YAP inhibits the expression of Zinc fingers and homeoboxes 2 (ZHX2), a VHL substrate and critical co-factor of NF-κB in ccRCC. Furthermore, YAP competes with ZHX2 for binding to the NF-κB subunit p65. Consequently, elevated nuclear YAP blocks the cooperativity between ZHX2 and the NF-κB subunit p65, leading to diminished NF-κB target gene expression. Pharmacological inhibition of Hippo kinase blocked NF-κB transcriptional program and suppressed ccRCC cancer cell growth, which can be rescued by overexpression of ZHX2 or p65. Our study uncovers a crosstalk between the Hippo and NF-κB/ZHX2 pathways and its involvement in ccRCC growth inhibition, suggesting that targeting the Hippo pathway may provide a therapeutical opportunity for ccRCC treatment.

## Introduction

Initially discovered in *Drosophila*, the Hippo tumor suppressor pathway is an evolutionarily conserved signaling pathway that controls organ size, tissue homeostasis, and cancer progression in different species (1–3). The Hippo pathway consists of a core kinase cascade in which the upstream kinases Hippo (Hpo)/MST1/2 phosphorylating and activating the downstream kinases LATS1/2 (4–8), resulting in the phosphorylation and inhibition of the pathway transcriptional effector YAP/TAZ (9–11). When the activity of Hippo kinase cascade is compromised, unphosphorylated YAP/TAZ translocates into the nucleus and forms a complex with the transcriptional enhanced associate domain (TEAD)-family of transcription factors to activate Hippo pathway target genes (11–13). Aberrant activation of YAP due to mutations in upstream components, gene amplification or fusion, or other unknown mechanisms promotes tumor progression in many types of cancer including hepatocellular carcinoma, lung adenocarcinoma, gastric cancer, colon cancer, mesothelioma, schwannomas, ependymomas, cervical squamous cell carcinoma, uveal melanomas, and esophageal squamous cell carcinoma (2, 3, 14). Consistent with these clinical observations, transgenic overexpression of YAP or knockout of upstream Hippo pathway components in mouse livers resulted in hepatomegaly, leading to the development of hepatocellular carcinoma (10, 15–20). Taken together, these observations have led to a prevalent view that Hippo signaling functions as a tumor suppressor pathway by blocking the oncogenic potential of YAP/TAZ. However, recent studies have revealed that YAP can function as a context-dependent tumor suppressor in several types of cancer including hematological cancers (21), estrogen receptor α (ERα) positive breast cancer (22, 23), androgen receptor (AR) positive prostate cancer (24), and VHL negative clear cell renal cell carcinoma (ccRCC) (25).

ccRCC makes up ∼80% of kidney cancer, which is among the top ten most diagnosed cancers around the world (26). ccRCC is responsible for most of the kidney cancer-associated death (27). The majority cases of ccRCC (>90%) are caused by loss-of-function mutations in the *Von Hippel-Lindau* (*VHL*) tumor suppressor gene leading to stabilization and activation of Hypoxia-inducible factor 2α (HIF-2α) (28). In addition to regulating HIF-2α, the VHL E3 ubiquitin ligase has other substrates that may also play important roles in the progression of ccRCC because ccRCC patient samples exhibited differential sensitivity to HIF-2α inhibitors (29–31). A genome-wide *in vitro* expression strategy identified Zinc fingers and homeoboxes 2 (ZHX2) as a VHL substrate (32). ZHX2 plays a pivotal role in ccRCC progression by activating the nuclear factor κB (NF-κB) pathway via interacting with the p65 subunit of NF-κB (32).

Our previous study revealed that inhibition of Hippo signaling pathway or transgenic activation of YAP blocked the HIF2 transcriptional program and ccRCC tumor growth (25). Here, we showed that YAP activation also inhibited NF-κB pathway in addition to HIF-2. Mechanistically, YAP acts in conjunction with TEAD to inhibit the expression of ZHX2 and its ability to bind p65, thereby blocking the cooperativity between ZHX2 and p65 required for NF-κB target gene expression and ccRCC growth. Pharmacological inhibition of Hippo/MST1/2 kinase activity inhibited p65/ZHX2 cooperativity, leading to diminished NF-κB target gene expression and inhibited ccRCC tumor growth, which can be alleviated by increasing the activity of ZHX2 or p65.

## Results

### YAP activation inhibits ccRCC cancer cell growth in 2D cultures

We have previously shown that activation of YAP, either by treatment with XMU-MP-1, a small molecule inhibitor of Hippo/MST1/2 kinase (33), or by overexpression of a constitutively active YAP (YAP-5SA) (9), inhibited ccRCC cell growth in both 3D cultures and Xenografts (25). Consistent with the observed cancer cell growth inhibition, both XMU-MP-1 and YAP5SA inhibited HIF-2α pathway activity that is required for ccRCC cell growth in 3D cultures or mice. Intriguingly, we found that XMU-MP-1 also inhibited cancer cell growth of multiple *VHL* mutant ccRCC cell lines including 786-O cells in 2D cultures where HIF-2α pathway activity was dispensable for their growth (Fig. 1A-C) (30, 34, 35), raising the possibility that XMU-MP-1 could inhibit ccRCC cell growth through a mechanism(s) other than blocking HIF-2α pathway activity.

**Figure 1.**
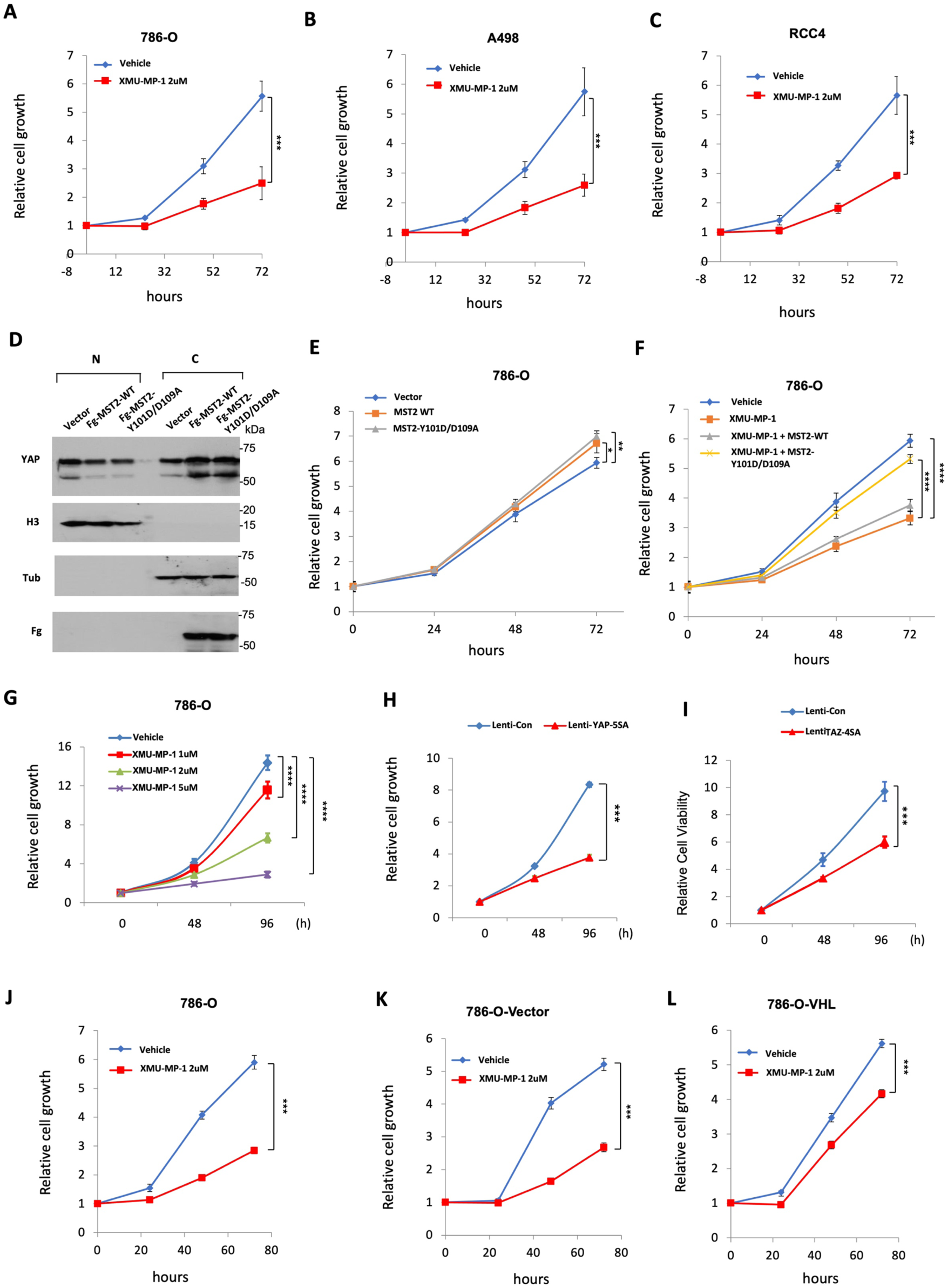
YAP activation inhibits ccRCC cancer cell growth in 2D cultures **A-C,** Growth of 786-O (**A**), A498 (**B**), and RCC4 (**C**) cells in 2D cultures treated with vehicle or 2 μM XMU-MP-1. **D,** Western blot analysis of YAP in nuclear (N) and cytoplasmic (C) fractions of 786-O cells expressing the indicated constructs. **E-F,** Growth of 786-O cells in 2D cultures expressing the indicated constructs in the absence (**E**) or presence (**F**) of XMU-MP-1 at 2 μM. **G,** Growth of 786-O cells in 2D cultures treated with increasing doses of XMU-MP-1. **H-I,** Growth of 786-O cells expressing the indicated lentiviral constructs in 2D cultures. **J-L,** Growth of 786-O cells (**J)** and 786-O cells expressing an empty vector (**K**) or VHL (**L**) and treated with vehicle or 2 μM XMU-MP-1. Data are ± SD. n=3 biological replicates. *P<0.05, **P<0.01, ***P<0.001, ****P<0.0001 (Student’s t-test for **A-C**, **H-L**; one-way ANOVA for **E-G)**.

To determine whether XMU-MP-1 inhibited ccRCC cell growth in 2D cultures through the Hippo pathway instead of an off-target effect, we generated Flag (Fg)-tagged wild type (Fg-MST2^WT^) and a mutant form of MST2 carrying Y101D and D109A double mutations (Fg-MST2^Y101D/D109A^). It has been demonstrated previously that MST2^Y101D/D109A^ exhibits normal kinase activity, but it no longer binds XMU-MP-1 and is insensitive to the drug inhibition (33). As expected, expression of either Fg-MST2^WT^ or Fg-MST2^Y101D/D109A^ increased the cytoplasmic level while decreased the nuclear level of YAP in 786-O cells, suggesting that exogenously expressed MST2 could increase the phosphorylation of YAP, leading to its cytoplasmic retention (Fig. 1D). Expression of either Fg-MST2^WT^ or Fg-MST2^Y101D/D109A^ in 786-O cells slightly increased cell growth (Fig. 1E); however, only Fg-MST2^Y101D/D109A^ significantly rescued XMU-MP-1-mediated growth inhibition (Fig. 1F). Hence, XMU-MP-1 inhibited 786-O cell growth mainly through an on-target effect, i.e., by inhibiting MST1/2 kinase activity to increase YAP nuclear localization.

To further investigate how perturbation of Hippo signaling affects ccRCC cell growth in 2D culture, we treated 786-O cells with different doses of XMU-MP-1 and found that XMU-MP-1 inhibited 786-O cell growth in a dose dependent manner (Fig. 1G). Furthermore, infection of 786-O cells with lentivirus expressing either the constitutively active YAP (YAP-5SA) or TAZ (TAZ-4SA) resulted in significant growth inhibition (Fig. 1H-I). To determine whether XMU-MP-1 inhibited ccRCC cell growth depending on the status of VHL, we obtained 786-O cells (786-O-VHL) with VHL-added back (36). Compared with original 786-O cell line or a control 786-O cell line expressing the empty vector (786-O-Vector), 786-O-VHL cells are less sensitive to XMU-MP-1-mediated growth inhibition (Fig. 1J-L), suggesting that XMU-MP-1 may act through another VHL target(s) to inhibit 786-O cell growth in 2D cultures.

### YAP activation inhibits the transcriptional program of NF-κB in ccRCC

A previous study identified ZHX2 as a VHL target upregulated in *VHL* mutant ccRCC tumors and showed that ZHX2 is essential for *VHL* mutant ccRCC cell growth *in vitro* and in *vivo* (32). Furthermore, this study demonstrated that ZHX2 promoted ccRCC tumor growth by promoting NF-κB pathway activity.

To determine whether XMU-MP-1 could inhibit ccRCC through the NF-κB pathway, we carried out RNAseq of 786-O cells treated with XMU-MP-1 or vehicle. Gene Ontology (GO) enrichment analysis of differentially expressed genes revealed that NF-κB signaling is among the top downregulated pathways in addition of HIF (Hypoxia) pathway (Fig.2A). Gene Set Enrichment Analysis (GSEA) revealed an enrichment of NF-κB pathway genes that were downregulated by XMU-MP-1(Fig. 2B). Further analysis of the RNAseq data indicated that XMU-MP-1 downregulated many NF-κB target genes that are coregulated by ZHX2 and p65, including *IL6*, *IL8* (*CXCL8*), *CCL2*, *TNF*, and *VCAM1* (Fig. 2C; Fig. S1)(32).

**Figure 2.**
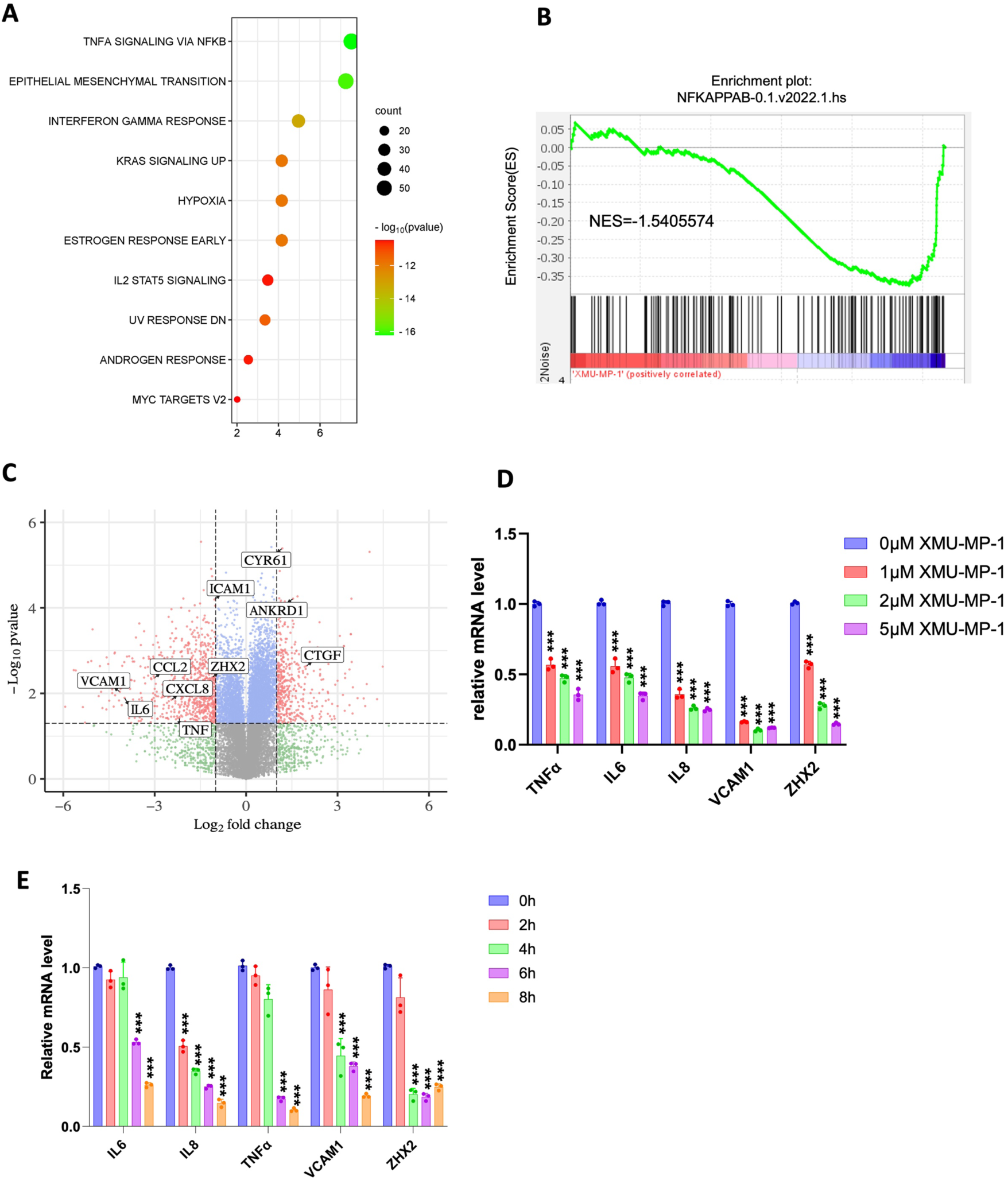
XMU-MP-1 inhibits NF-κB transcriptome in ccRCC **A,** Top 10 signaling pathways downregulated in 786-O cells treated with XMU-MP-1. **B,** Gene set enrichment analysis (GSEA) shows downregulation of NF-κB target genes in 786-O cells treated with XMU-MP-1. **C,** Volcano plot shows the opposite effects of XMU-MP-1 treatment on NF-κB target genes and the Hippo pathway signature in 786-O cells treated with XMU-MP-1. **D,** Relative mRNA levels of the indicated NF-κB target genes and *ZHX2* in 786-O cells treated with the indicated concentrations of XMU-MP-1. **E,** Relative mRNA levels of the indicated NF-κB target genes and *ZHX2* in 786-O cells treated with XMU-MP-1 for 0, 2, 4, 6, or 8 hours. Data in **D** and **E** are ± SD. n=3 biological replicates. ***P<0.001 (one-way ANOVA).

To confirm that Hippo pathway regulates NFκB signaling in ccRCC, we carried out RT-qPCR experiments to measure the change in NFκB target gene expression in ccRCC upon treatment with XMU-MP-1 at different doses or at a fixed concentration for increasing amount of time. As shown in Fig. 2D, high dose of XMU-MP-1 resulted in a more dramatic downregulation of NFκB target gene expression. Similarly, increasing the duration of XMU-MP-1 treatment progressively decreased the expression of multiple NF-κB target genes without affecting the expression of p65 protein level (Fig. 2E; Fig. S2). These results suggest that XMU-MP-1 inhibits NFκB target gene expression in a dose- and time-dependent manner and that this downregulation is not due to a change in the abundance of p65, a major subunit of NFκB in ccRCC (32). Activation of YAP by the constitutively active YAP5SA resulted in inhibition of NF-κB target gene expression (Fig. 3A, B). On the contrary, knockdown of both YAP and TAZ using a previously validate siRNA (24, 25), increased NFκB target gene expression (Fig. 3C, D). These results suggest that YAP negatively regulates NFκB signaling activity.

**Figure 3.**
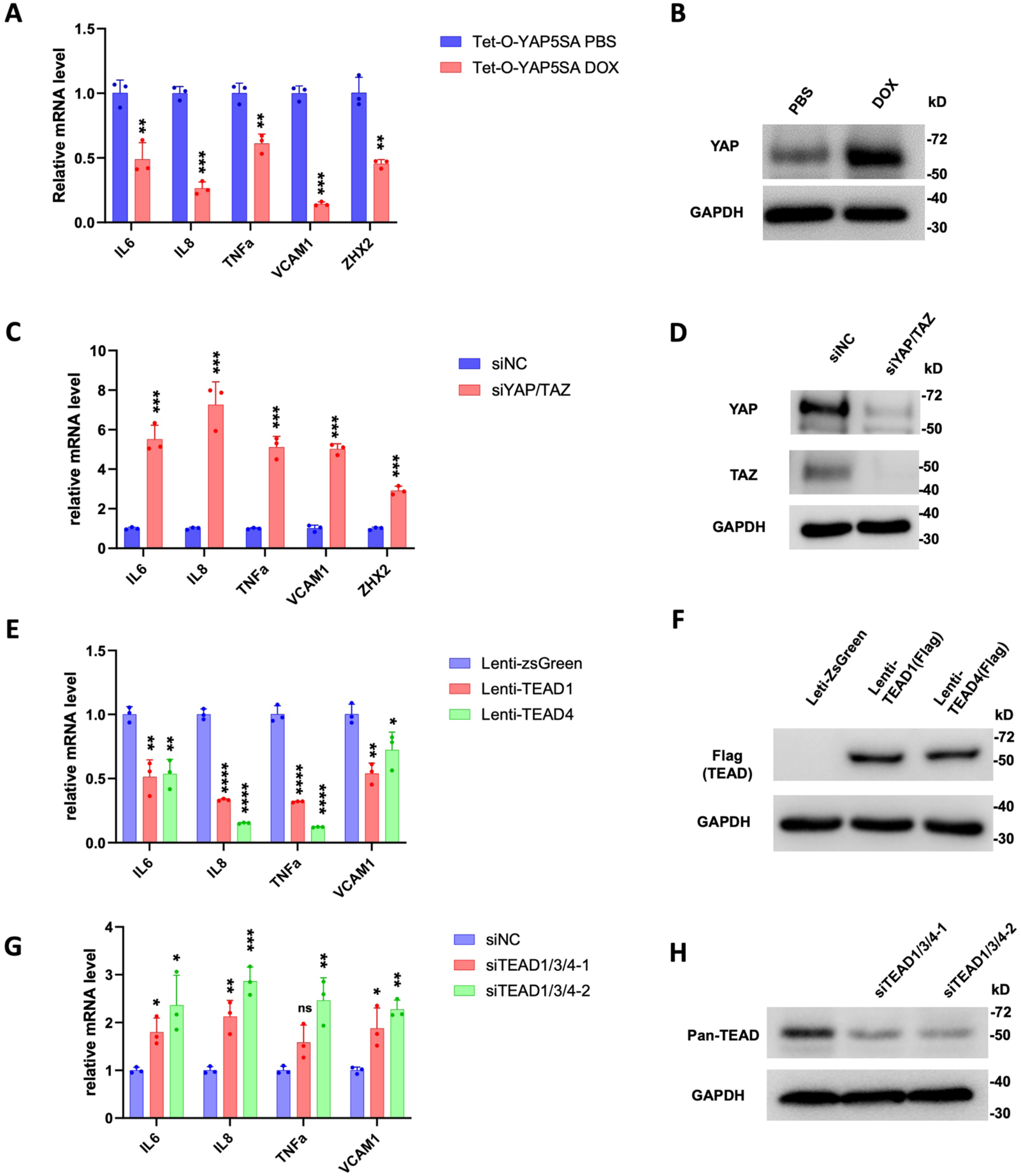
YAPTEAD inhibits NF-κB target gene expression in ccRCC **A,** Relative mRNA levels of the indicated NF-κB target genes and *ZHX2* in 786-O cells expressing Tet-O-YAP5SA and treated with PBS or Doxycycline (Dox). **B,** Western blot to examine YAP protein level in 786-O cells expressing Tet-O-YAP5SA and treated with PBS or Dox. **C,** Relative mRNA levels of the indicated NF-κB target genes and *ZHX2* in 786-O cells treated with control siRNA (siNC) or a siRNA targeting both YAP and TAZ (siYAP/TAZ). **D,** Western blot to examine YAP/TAZ protein levels in 786-O cells treated with siNC or siYAP/TAZ. **E,** Relative mRNA levels of the indicated NF-κB target genes in 786-O cells infected with control, TEAD1-Flag or TEAD4-Flag lentivirus. **F,** Western blot to examine TEAD1/TEAD4-Falg level in 786-O cells infected with control, TEAD1-Flag or TEAD4-Flag lentivirus. **G,** Relative mRNA levels of the indicated NF-κB target genes in 786-O cells treated with control siRNA or siRNAs targeting TEAD1/3/4. **H,** Western blot to examine TEAD levels in genes in 786-O cells treated with control siRNA or siRNAs targeting TEAD1/3/4. Data are ± SD. n=3 biological replicates. *P<0.05, **P<0.01, ***P<0.001, ****P<0.0001 (Student’s t-test for **A**, **C**; one-way ANOVA for **E**, **G**). ns: not significant.

YAP regulates gene expression through the TEAD family of transcription factors including TEAD1-4 (37). RNAseq analysis revealed that TEAD1 and TEAD4 were more abundantly expressed than TEAD2 and TEAD3 in 786-O cells (25). We found that overexpression of either TEAD1 or TEAD4 downregulated NF-κB target gene expression (Fig. 3E, F). On the other hand, knockdown of TEAD1/3/4 using previously validate siRNAs that targets shared sequences among these TEADs led to increased expression of NF-κB target genes (Fig. 3G, H). These results suggest that YAP acts in conjunction with TEAD to inhibit NFκB target gene expression.

### YAP inhibits *ZHX2* expression

Analysis of the RNAseq data revealed that XMU-MP-1 treatment of 786-O cells also downregulated the expression of *ZHX2*. By RT-qPCR, we confirmed that XMU-MP-1 decreased the expression of ZHX2 in 786-O cells (Fig. 2D, E). Expression of YAP-5SA also downregulated whereas knockdown of YAP/TAZ increased *ZHX2* expression in 786-O cells (Fig. 3A, C), suggesting that YAP inhibits *ZHX2* expression.

Because ZHX2 is a critical cofactor for p65 in the regulation of NFκB target gene expression, YAP-mediated downregulation of ZHX2 could explain why XMU-MP-1 and YAP-5SA inhibit the expression of NFκB target genes. However, in a time course experiment, we found that XMU-MP-1 inhibited NFκB target genes as well as *ZHX2* within 8 hours whereas ZHX2 protein levels were noticeably downregulated only after 16 hours treatment (Fig. S3A-C). Whereas transcriptional downregulation of *ZHX2* may contribute to a long-term shutdown of NFκB target gene expression by YAP activation, the immediate response of NF-κB target genes to XMU-MP-1 treatment is likely due to a different mechanism(s).

### YAP blocks the interaction between ZHX2 and p65

Because ZHX2 promotes NFκB target gene expression by physically interacting with p65, we carried out co-immunoprecipitation (Co-IP) experiments to determine whether YAP inhibits NFκB pathway activity by inhibiting p65/ZHX2 association. Using antibodies against endogenous proteins, we found that p65 formed a complex with ZHX2 as well as YAP and TEAD4 (Fig. 4A). Interestingly, XMU-MP-1 decreased the amount of ZHX2 associated with p65 while simultaneously increased the amount of YAP and TEAD4 bound to p65 (Fig. 4A), suggesting that YAP-TEAD may compete with ZHX2 for binding to p65.

**Figure 4.**
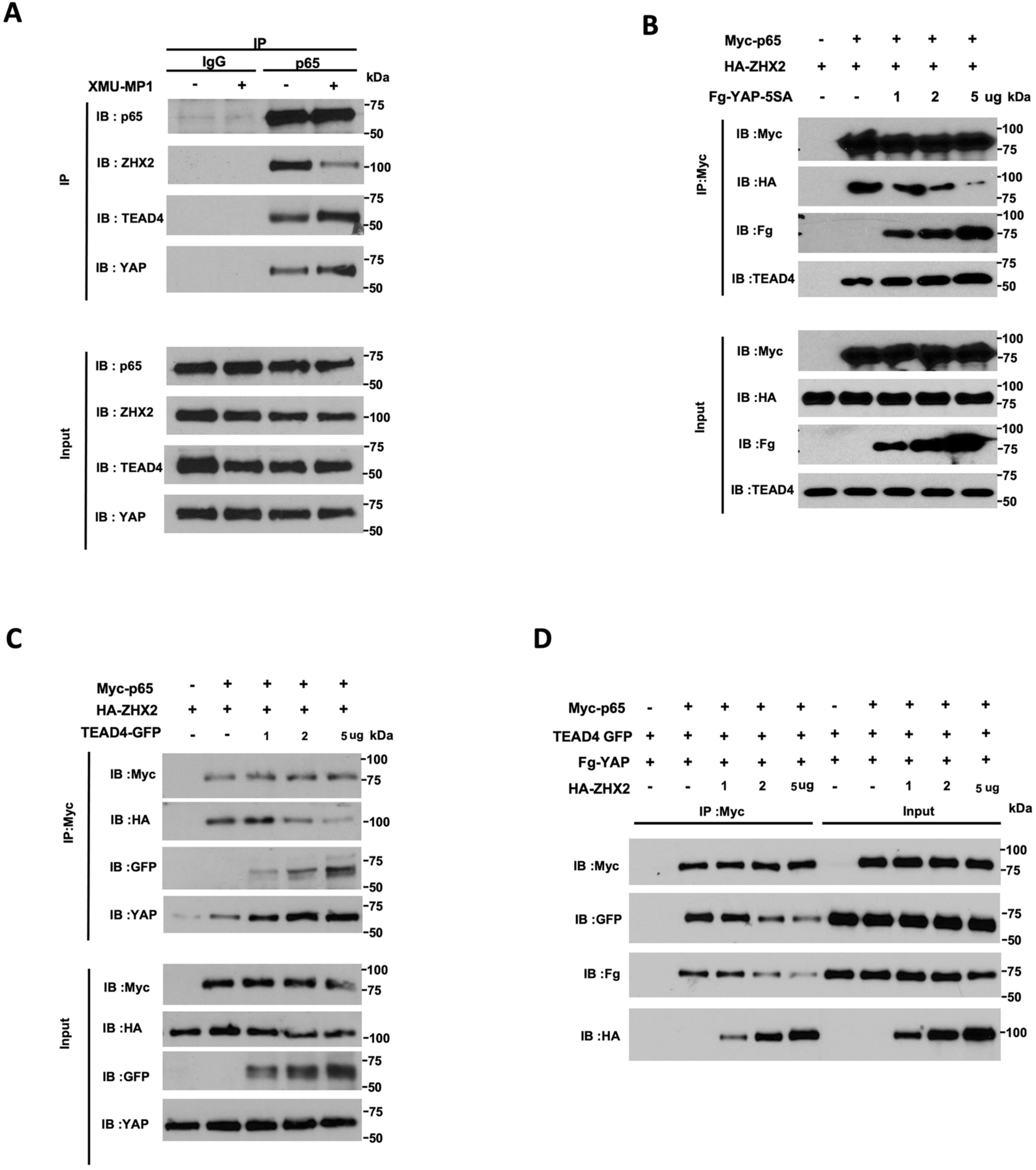
YAP/TEAD competes with ZHX2 for binding to p65 **A,** Western blot analysis of endogenous p65, ZHX2, YAP and TEAD4 immunoprecipitated with control (IgG) or p65 antibody from cell extracts of 786-O cells grown without or with 2 μM XMU-MP-1 for 6 hours. **B-C,** HEK293 cells were transfected with fixed amounts of Myc-p65 and HA-ZHX2 and increasing amounts of Fg-YAP-5SA (**B**) or TEAD4-GFP (**C**). Cell extracts were immunoprecipitated with an anti-Myc antibody, followed by western blot analysis with antibodies against the indicated tags, endogenous TEAD4 (**B**) or YAP (**C**). **D,** HEK293 cells were transfected with fixed amounts of Myc-p65, Fg-YAP and TEAD4-GFP and increasing amounts of HA-ZHX2. Cell extracts were immunoprecipitated with the anti-Myc antibody, followed by western blot analysis with antibodies against the indicated tags.

To further characterize the competition between YAP-TEAD and ZHX2 for p65 binding, HEK293 cells were transfected with fixed amounts of Myc-tagged p65 (Myc-p65) and HA-tagged ZHX2 (HA-ZHX2) and increasing amounts of Flag-tagged YAP-5SA (Fg-YAP-5SA) or GFP-tagged TEAD4 (TEAD4-GFP), followed by Co-IP and western blot analysis. As shown in Fig. 4B, increasing the expression level of Fg-YAP-5SA progressively increased the amount of p65/Fg-YAP-5SA complex with concomitant decrease in the amount of p65-ZHX2 complex. Furthermore, increasing the amount of Fg-YAP-5SA also resulted in an increase in the amount endogenous TEAD4 associated with Myc-p65. Similarly, increasing the amount of TEAD4-GFP led to a decrease in the amount of HA-ZHX2 associated with Myc-p65 but an increase in the amounts of TEAD4--GFP and endogenous YAP associated with Myc-p65 (Fig. 4C). These results suggest that binding of YAP and TEAD to p65 disrupt the formation of the p65-ZHX2 complex. In a reciprocal experiment, HEK293 cells were transfected with fixed amounts of Myc-p65, Fg-YAP5SA and TEAD4-GFP, and increasing amount of HA-ZHX2. As shown in Fig. 4D, increasing the expression level of HA-ZHX2 progressively increased the amount of p65-ZHX2 complex with concomitant decrease in the amounts of Fg-YAP5SA and TEAD4-GFP associated with Myc-p65. Taken together, these results demonstrate that YAP-TEAD and ZHX2 compete for binding to p65.

### YAP impedes binding of p65 and ZHX2 to NF-κB target promoters

ZHX2 and p65 co-occupy on the promoter/enhancer regions of many NFκB target genes and ZHX2/p65 interaction facilitates their promoter/enhancer occupancy to NF-κB target genes as knockdown of ZHX2 impaired p65 binding to *IL6* and *IKBKE* promoters, and vice versa (32). Therefore, we tested whether YAP activation by Hippo pathway inhibition reduces the binding of p65/ZHX2 to the promoter regions of their coregulated genes by carrying ChIP-qPCR experiments. As shown in Fig. 5, after 786-O cells were treated with XMU-MP-1, both ZHX2 and p65 exhibited decreased chromatin association at multiple NF-κB target promoters co-bound by p65 and ZHX2, including *IL6*, *IL8*, and *CCL2* (Fig. 5A-C). Interestingly, p65 and ZHX2 co-occupied the promoter region of *ZHX2*, which was reduced by XMU-MP-1 treatment (Fig. 5A-C), suggesting that *ZHX2* is an autoregulated NF-κB target gene and that YAP inhibits *ZHX2* expression by disrupting p65/ZHX2 interaction.

**Figure 5.**
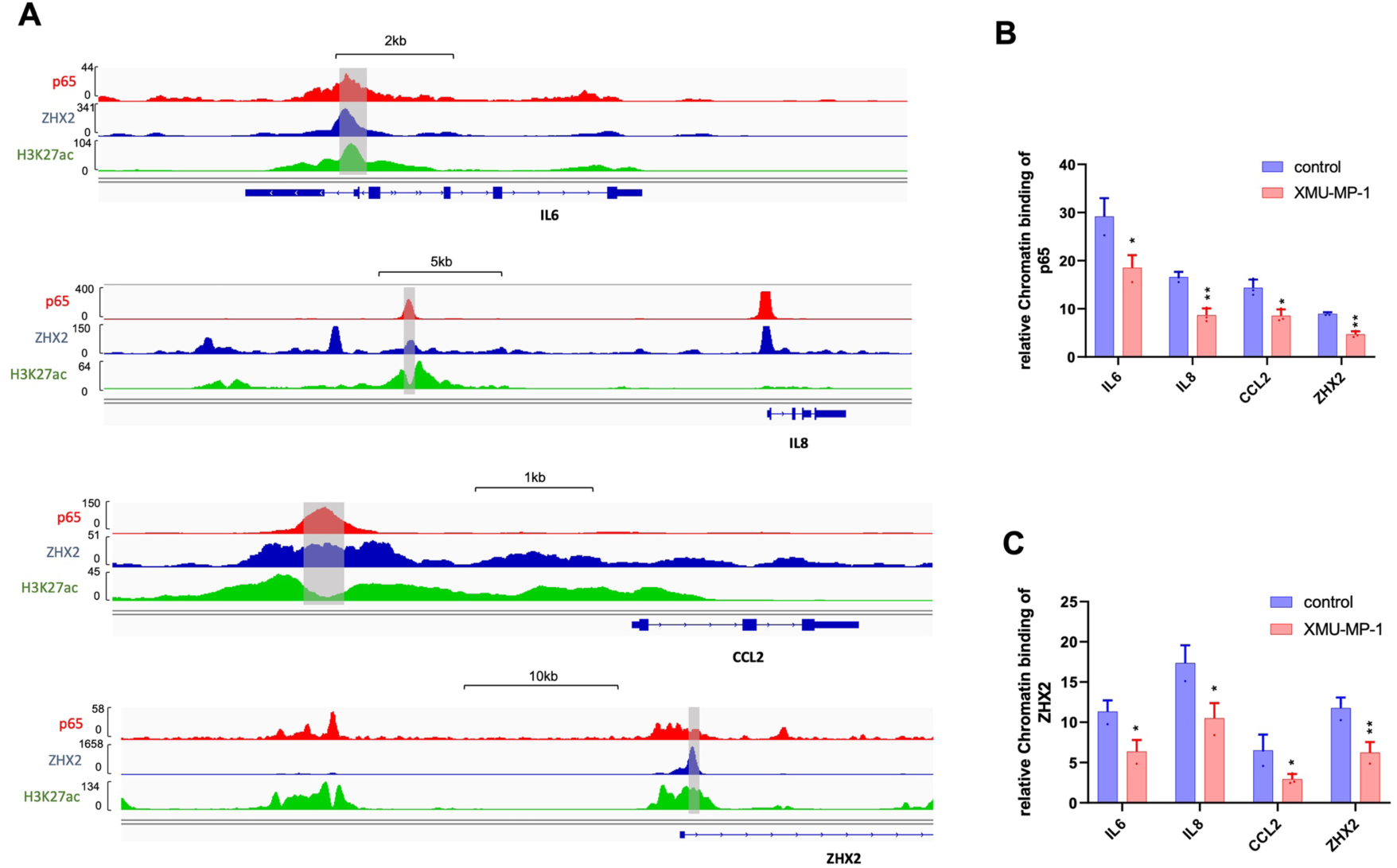
Hippo pathway inhibition impairs occupancy of p65 and ZHX2 on NF-κB target promoters **A,** Analysis of a published ChIP seq data indicates that p65 and ZHX2 co-bind sites in the promoter/enhancer regions of *IL6*, *IL8*, *CCL2* and *ZHX2* that are also enriched for H3K27ac. **B-C,** ChIP qPCR experiments show that treating 786-O cells with the Hippo pathway inhibitor XMU-MP-1 decreased the binding of p65 (**B**) and ZHX2 (**C**) to the promoter/enhancer regions (shaded regions in **A**) of *IL6*, *IL8*, *CCL2* and *ZHX2*. Data in are ± SD. n=2 biological replicates. *P<0.05,**P<0.01 (Student’s t-test).

### ZHX2/p65 overexpression rescued ccRCC growth inhibited by XMU-MP-1

If Yap activation inhibits ccRCC cancer cell growth in 2D cultures by impeding p65/ZHX2 cooperativity, one would predict that increasing the level of either ZHX2 or p65 may rescue ccRCC cancer cell growth inhibited by XMU-MP-1. To test this possibility, we overexpressed ZHX2 or p65 in 786-O cells via lentiviral infection and measured cancer cell growth in 2D cultures as well as NF-κB target gene expression in the absence or presence of XMU-MP-1 treatment. As expected, overexpression of either ZHX2 or p65 increased NF-κB target gene expression and promoted ccRCC cell growth in the absence of drug treatment (Fig. 6A-F).

**Figure 6.**
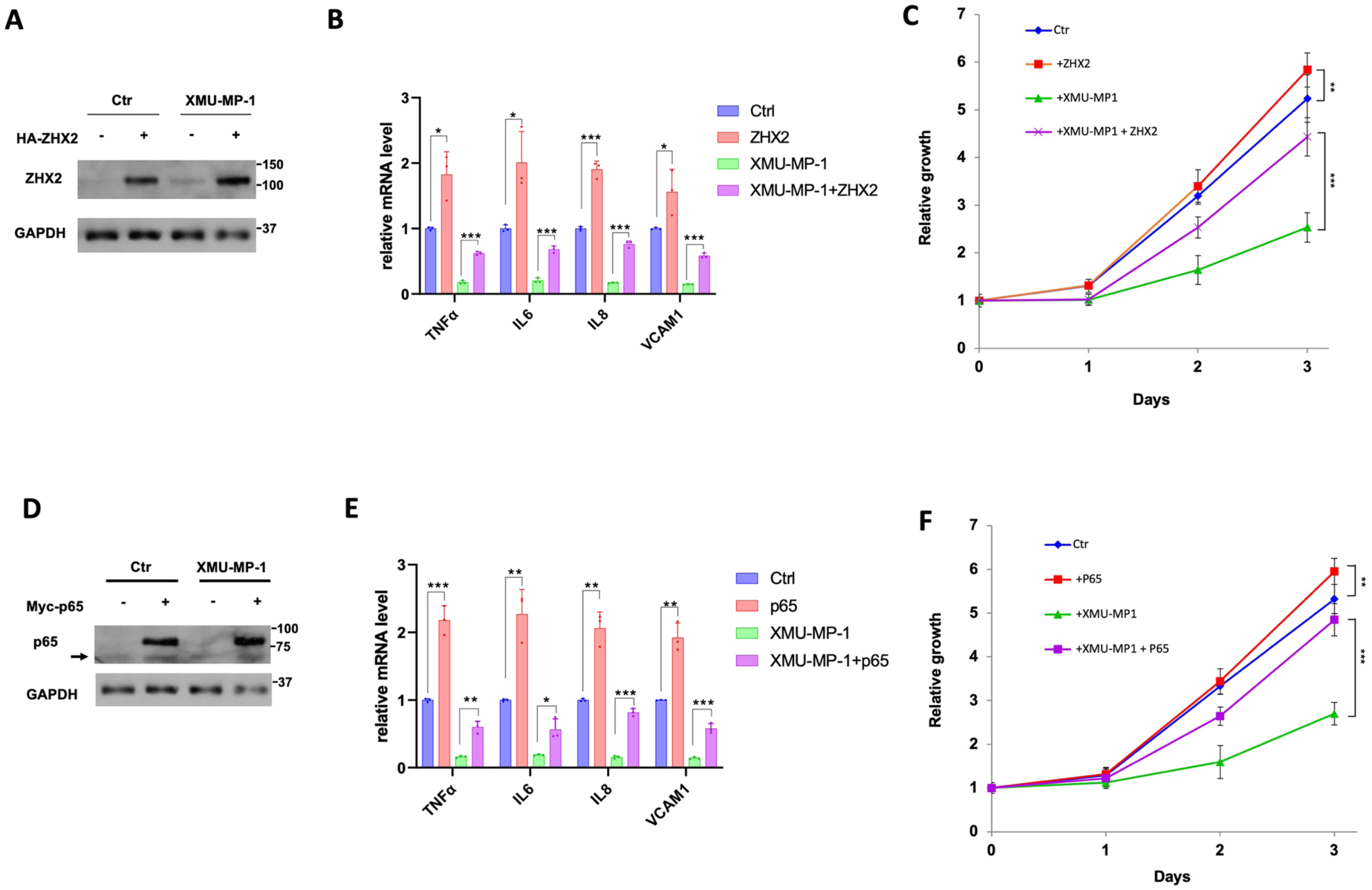
Overexpression of p65/ZHX2 rescues ccRCC growth inhibited by XMU-MP-1. **A-B,** Western blot analysis of ZHX2 (**A**) and RT-qPCR analysis of the indicated NF-κB target gene expression (**B**) in 786-O cells infected with control or ZHX2 expressing lentivirus and treated with vehicles or 2 μM XMU-MP-1. Data in are ± SD. n=3 biological replicates. *P<0.05,***P<0.001 (Student’s t-test). **C,** Growth of 786-O cells infected with control or ZHX2 expressing lentivirus and treated with vehicles or 2 μM XMU-MP-1. Data in are ± SD. n=3 biological replicates. **P<0.01, ***P<0.001 (Student’s t-test). **D-E**, Western blot analysis of p65 (**D**) and RT-qPCR analysis of the indicated NF-κB target gene expression (**E**) in 786-O cells infected with control or p65 expressing lentivirus and treated with vehicles or 2 μM XMU-MP-1. Arrow in **D** indicates endogenous p65. Data in are ± SD. n=3 biological replicates. *P<0.05,**P<0.01, ***P<0.001 (Student’s t-test). **F,** Growth of 786-O cells infected with control or p65 expressing lentivirus and treated with vehicles or 2 μM XMU-MP-1. Data in are ± SD. n=3 biological replicates.**P<0.01, ***P<0.001 (Student’s t-test).

Importantly, ZHX2 or p65 overexpression significantly rescued ccRCC cell growth and NF-κB target gene expression after XMU-MP-1 treatment (Fig. 6A-F), lending further support that YAP inhibits ccRCC cancer cell growth by opposing the cooperativity between p65 and ZHX2.

## Discussion

Despite the prevailing view that Hippo signaling inhibits tumor growth by blocking the oncogenic potential of YAP/TAZ (2, 14), recent studies revealed that YAP can act as a context-dependent tumor suppressor in several types of cancer (21–25, 38–40). Our previous study uncovered a noncanonical Hippo signaling mechanism in ccRCC whereby TEAD functions as a critical cofactor for the ccRCC oncogenic driver HIF-2α whereas nuclear YAP inhibits HIF-2α signaling by competing with HIF-2α for TEAD (25). Analogous mechanisms have also been proposed to account for the tumor suppressor function of YAP in ERα^+^ breast cancer and AR^+^ prostate cancer (23, 24). In the current study, we uncovered yet another mechanism by which YAP inhibits cancer cell growth, i.e., through inhibiting the NFκB signaling pathway.

Previous studies showed that the growth of *VHL^-/-^* ccRCC cancer cells in 2D cultures was independent of HIF-2α activity (30, 34, 35). Our observation that XMU-MP-1 could inhibit *VHL^-/-^* ccRCC growth not only in soft agar (3D cultures) but also in 2D cultures, a condition in which HIF-2α is dispensable, implied that YAP may inhibit *VHL^-/-^*ccRCC growth through additional mechanism(s). Furthermore, such inhibitory mechanism depends on the status of VHL as restoring VHL activity in 786-O cells blunted the inhibitory effect of XMU-MP-1, suggesting that YAP may inhibit 786-O cell growth through another VHL substrate(s). The recent finding that ZHX2 is a VHL substrate that promotes ccRCC cancer progression by forming a complex with p65 to regulate NF-κB target genes let us to speculate that YAP may target the p65-ZHX2 signaling axis. Indeed, we found that treatment of 786-O cells with XMU-MP-1 downregulated many NF-κB target genes coregulated by p65 and ZHX2 (Fig. S1). XMU-MP-1 treatment disrupted the interaction between p65 and ZHX2 while increasing the interaction between p65 and YAP-TEAD. Co-IP experiments in a heterologous system further demonstrate the competition between YAP-TEAD and ZHX2 for p65 binding. Increasing the level of ZHX2 or p65 rescued ccRCC growth inhibited by XMU-MP-1. Based on these and other observations, we propose that YAP-TEAD inhibits NF-κB signaling in ccRCC by opposing p65/ZHx2 cooperativity, which contributes to ccRCC growth inhibition (Fig. 7).

**Figure 7.**
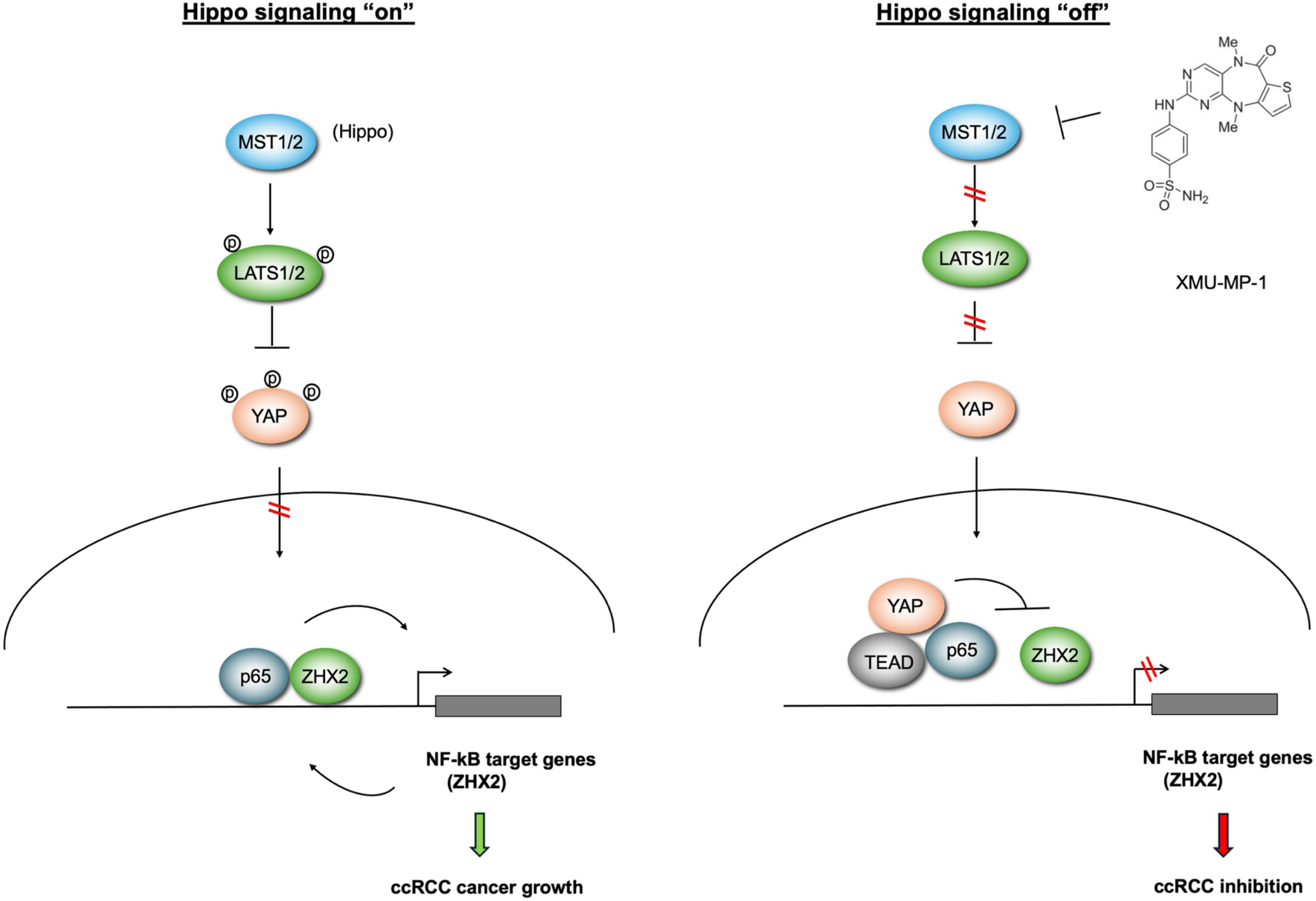
Hippo pathway inhibition impedes ccRCC growth by interfering with p65/ZHX2 cooperativity When Hippo signaling is “on”, YAP is phosphorylated and excluded from the nucleus, allowing p65 and ZHX2 to cooperatively activate NF-κB target gene expression that drives ccRCC growth. When Hippo signaling is “off”, YAP enters the nucleus and forms a complex with TEAD. YAP/TEAD impedes p65/ZHX2 cooperativity by disrupting their physical interaction, leading to downregulation of NF-κB target gene expression and ccRCC growth inhibition.

A recent study suggested that TEAD is a co-activator of p65, and that YAP inhibit NF-κB signaling by competing with p65 for binding to TEAD (41). However, our CoIP experiments clearly showed that binding of YAP to p65 did not disrupt but rather increasing p65-TEAD interaction (Fig. 4). Consistently, binding of TEAD to p65 also increased the association between p65 and YAP, suggesting that YAP and TEAD act cooperatively to regulate NF-κB signaling, which is different from the mechanisms by which YAP inhibits ERα, AR, and HIF2α (23–25).

ccRCC is responsible for most of the kidney cancer-associated death. Therapies targeting HIF-2α downstream effectors such as VEGF are the standard of care; however, drug resistance occurs in most patients (31, 42). HIF-2α inhibitors have recently entered the clinic; however, ccRCC patient samples exhibited differential sensitivity to HIF-2α inhibitors and HIF-2α mutants could render drug resistance (29, 43). Our findings that Hippo pathway inhibition or YAP activation not only inhibits HIF2α but also NF-κB signaling in ccRCC open a possibility for developing novel therapeutics to treat ccRCC.

### Experimental procedures

#### DNA constructs

Myc-p65, HA-ZHX2, TEAD4-GFP, VHL, TAZ4SA and Flag-YAP were amplified by PCR and the products were sub-cloned to pLVX-zsGreen vector. The Flag-MST2-wt were amplified by PCR. To generate Flag-MST2-dm, Y101F/D109A double mutation was amplified by PCR-based mutagenesis from Flag-MST2, and the product was subcloned to pLVX-ZsGreen vector. The YAP-WT and YAP-5SA constructs were described previously (44). YAP-WT or YAP-5SA were cloned into the pLVX-IRES-ZsGreen vector. GFP and YAP5SA coding sequences were subclone into pTet-O-Ngn2-puro (Addgene plasmid, Cat. No 52047) to generate Tet-O-GFP and Tet-O-YAP5SA. pLenti CMV rtTA3 Hygro (Addgene plasmid, Cat. No. 26730) was used to generate Dox-inducible cell lines.

#### Cell cultures

786-O, A498 and RCC4 are maintained with RPMI-1640 (Gibco, Cat. No. 42401018) supplemented with 2 mM L-glutamine (Gibco, Cat. No. 25030081) and 10% FBS. The HEK293A and HEK293T cells are culture with high glucose Dulbecco’s Modified Eagle’s Medium that contains 4.5 g/L glucose and 4 mM L-glutamine (DMEM, Gibco, Cat. No. 11965092) supplemented with 10% Fetal Bovine Serum (FBS, Gibco, Cat. No. 16000044).

#### Cell proliferation assay

∼3,000 ccRCC cells were seeded into 96-well plates. After cells were attached to the plates, they were treated with XMU-MP-1 for the indicated concentrations or periods of time. For 786-O cells expressing different constructs, ∼3,000 cells were seeded into 96-well plates. The relative cell viability was measured at the indicated time points. Cell numbers were determined using the WST-1 cell proliferation reagent (Sigma Aldrich, Cat. No. 5015944001).

#### Immunoblot analysis

Cells were harvested and lysed with lysis buffer containing 1M Tris pH8.0, 5M NaCl, 1M NaF, 0.1M Na_3_VO_4_, 1% NP-40, 10% glycerol, and 0.5M EDTA (pH 8.0). Proteins were separated by electrophoresis on SDS-polyacrylamide gel electrophoresis (PAGE) and electro-transferred to PVDF membrane. Membranes were washed with TBST and incubated with primary antibodies for 2 hours. And then the membranes were washed for three times with TBST and incubated with second antibodies for 2 hours, after washed for three times with TBST, the membranes were probed with ECL system (Cytiva, Cat. No. RPN2105). The antibodies used in this study were listed here: rabbit anti-p65 (Cell Signaling Technology Cat.No,8242); rabbit anti-ZHX2 (Genetex, Cat.No. 112232); mouse anti-H3 (Santa Cruz, Cat. No. sc-517576); mouse anti-tubulin (Santa Cruz, Cat. No. sc-5288); mouse anti-GAPDH (Santa Cruz, Cat. No. sc-47724); mouse anti-YAP (Santa Cruz, Cat. No. sc-101199); mouse anti-TEAD4 (Santa Cruz, Cat. No. sc-101184); rabbit anti-HA (COVANCE, Cat. No. MMS-101R); mouse anti-Myc (Santa Cruz, Cat.No.SC-40); anti-Flag (Sigma, Cat. No. F3165); anti-GFP (Abcam, Cat. No. ab290). Peroxidase-Conjugated AffiniPure goat anti-mouse IgG (Jackson ImmunoResearch, Cat.No. 115-035-003) or goat anti-rabbit IgG (Jackson ImmunoResearch, Cat.No.111-035-144). Chemiluminescent signals were visualized with ECL system (Cytiva, Cat. No. RPN2105).

#### Co-immunoprecipitation assay

Immunoprecipitation was performed according to standard protocol. 786-O cell lysates were incubated with antibodies or mouse IgG for overnight at 4℃, followed by immobilization and precipitation with Protein A resin (Thermo Scientific Cat.No53139). The bound proteins were analyzed by western blot. For overexpression experiments, HEK293A cells were transfected with 5μg myc-p65, HA-ZHX2, Flag-YAP and GFP-TEAD4 plasmids in 10 cm dishes. Cell lysates were incubated with antibodies against epitope tags for overnight, followed by immobilization and precipitation with Protein A resin. The bound proteins were analyzed by immunoblot assay. Antibodies used for IP experiments: anti-p65 (Cell Signaling Technology Cat.No,8242), anti-ZHX2 (Genetex, Cat.No. 112232), anti-YAP (Santa Cruz, Cat. No. SC-101199), anti-TEAD4 (Santa Cruz, Cat. No. sc-101184), anti-Flag (Sigma, Cat.No. F3165), anti-Myc (Santa Cruz, Cat.No.SC-40), anti-GFP (Abcam, Cat. No. ab290), or mouse IgG (Santa Cruz, Cat.No. SC-3881).

#### RNA interference

For RNAi experiments in renal cancer cells, the siRNAs were acquired from the Sigma-Aldrich. The RNAi MAX reagent (Invitrogen Cat. No. 13778150) was used for the transfection of siRNA according the manuscription. Knockdown efficiency was validated by RT-PCR and/or immunoblotting. The sequences for YAP/TAZ silencing were: 5’-UGU GGA UGA GAU GGA UA CA-3’ and 5’-UGT AUC CAU CUC AUC CAC A-3’. The sequences for TEAD1/3/4 silencing were: 5’-AUG AUC AAC UUCA UCC ACA AG-3’ and 5’ GAU CAA CUU CAU CCA CAA GCU-3’. The sequences for negative control were: 5’-UUC UCC GAA CGU GUC ACG U-3’.

#### Virus infection and transient transfection

For packaging lentivirus, HEK293T cells were transfected with the expression vectors and package vectors (psPAX2 and pMD2.G) by PolyJet (SignaGen laboratories, Cat. No. SL100688). After 48 hours, the supernatants of the medium were collected and filtered with 0.45 μm filter. The supernatant containing virus was stored in 4℃ for cell infection. The ccRCC cells were cultured in fresh media and subsequently infected with lentivirus overnight together with Polybrene (Sigma, Cat. No. H9268). Hygromycin B (Sigma, Cat. No. 10843555001) and Puromycin (Sigma, Cat. No. P9620) were used for infected cells selection according to the resistance of the vectors.

#### RNA extraction and RT-qPCR analysis

We extracted the total RNA by RNeasy plus mini kits according to the protocol (Qiagen, Cat. No. 74106). After RNA extraction, the RNA was subjected to reverse transcription PCR for cDNA synthesis according to the RT-PCR kit (Applied Biosystems, Cat. No. 4368814). The relative gene expression was measured according to 2^-ΔΔCT^ methods. The house keeping gene 36B4 was used for internal control. The Primer sequences were: 36B4, F: GGC GAC CTG GAA GTC CAA CT; R: CCA TCA GCA CCA CAG CCT TC;IL6, F: AGA CAG CCA CTC ACC TCT TCA G, R: TTC TGC CAG TGC CTT TTG CTG; IL8, F: GAG AGT GAT TGA GAG TGG ACC AC, R: CAC AAC CCT CTG CAC CCG ATT T; TNF, F: CTC TTC TGC CTG CTG CAC TTT G; R: ATG GGC TAC AGG CTT GTC ACT C; CCL2, F: AGA ATC ACC AGC AGC AAG TGT CC, R: TCC TGA ACC CAC TTC TGC TTG G; VCAM1, F: GAT TCT GTG CCC ACA GTA AGG C, R: TGG TCA CAG AGC CAC CTT CTT G; ZHX2, F: ACA CGG GAC CGA TAT GAT GC, R: TTG GAG GGG GAT AAG GAG GG.

#### ChIP qPCR

ChIP (Chromatin Immuno-precipitation) assay was performed as previously study description (45), In brief, cells were cross-linked using 2 mM DSG crosslinker (CovaChem, Cat. No.13301) at room temperature for 1 h, followed by secondary fixation with 1% formaldehyde (Pierce, Cat. No. 28908) for 10 min and quenched by glycine. Subsequently, cells were washed with cold PBS and subject to cell lysis. The cell extracts were sonicated by Bioruptor. After centrifuge, the supernatants were incubated with prepared p65/ZHX2/TEAD4 antibody-Dynabeads for 1 hour room temperature and another 1 hour at 4℃. The slurries were washed in wash buffer for 5 times and de-cross-linked ChIP in elution buffer at 65℃ overnight. The enriched DNA was extracted via DNA extraction kits (Qiagen, Cat. No. 28106) and subject to quantitative PCR analysis. The Primer sequences for ChIP-qPCR were: IL6, F: CTT CGT GCA TGA CTT CAG CTT T, R: AGG GGG AAA AGT GCA GCT TAG; IL8, F: TTC CAC ACA TGG TCA AGG GG, R: CCT TCT CCA GGC TCC ATT CA; ZHX2, F: ATT GCA CGG AGA CGG TTT GG, R: ACG GAC TCG GTG GAA TTT CT; CCL2, F: TTG TGC CGA GAT GTT CCC AG, R: TGG CGT GAG AGA AGT GAG TG. The antibodies used in ChIP-qPCR were anti-p65 (Cell Signaling Technology Cat.No,8242), anti-TEAD4 (Santa Cruz, Cat. No. SC-101184) and anti-ZHX2 (Genetex, Cat.No. 112232). Anti-rabbit IgG dynabeads (Invitrogen, Cat: 11204D) and anti-mouse IgG dynabeads (Invitrogen, Cat. No. 11031) were used to bind antibodies.

#### RNA sequencing and data analysis

The global gene expression analysis (Vehicle vs XMU-MP-1 treated groups) was based on RNA sequencing platform from BGI (Beijing Genomic Institute). Cellular RNA was extracted using Qiagen RNA extraction kit (Qiagen; Cat: 74104) according to the manufacturer’s instructions.

The cellular RNA was sent to BGI Genomics (https://www.bgi.com) for RNA sequencing. RNA was quality-accessed with an Agilent 2100 Bioanalyzer (Agilent RNA 6000 Nano Kit) with RNA integrity number above 9 for library construction. The total RNA was used for library construction according to the protocol of BGISEQ-500 platform. The libraries were sequenced using BGISEQ-500 platform. Then the FASTQ sequencing files were aligned to the hg19 human genome using STAR aligner with uniquely mapped reads kept for further analysis. Differential expression was analyzed using DESeq2 with default parameters. The RNA sequence data are deposited in the Gene Expression Omnibus (GEO) database (Assessing number: GSE197468). For gene set enrichment analysis of RNA-seq data, gene sets of HIF2A activated target genes was used and downloaded from Molecular Signatures Database v7.4, GSEA was implemented using the GSEA 4.1.0 software, with default parameters. Volcano plot of DEGs (Threshold P<0.01 and fold change>2) was generated using the OmicStudio tools (https://www.omicstudio.cn/tool).

## Statistics and reproducibility

All experiments were performed at least three independent times unless noted. Student’s t-test or one-way Anova was used for comparisons. P-value of < 0.05 was significant. Error bars on the graphs were presented as the s.d.

## Data availability

All data are contained within the manuscript.

## Supporting information

This article contains supporting information.

## Acknowledgements

We thank Drs. James Brugarolas and Qing Zhang for providing ccRCC cell lines.

## Author contributions

X.L., Y.C., Y.H.,S.Z., performed the experiments. X.L., Y.C., Y.H., M.Z, Y.L., Y.Y, S.Z., J.J. analyzed the data, X.L., S.Z., and J.J. designed the experiments. X.L. and J.J. wrote the manuscript.

## Funding and additional information

The work was supported by grants from NIH to JJ (R35GM118063, P50CA196516) and YY (R01CA222571), and Welch foundation (I-1603) to JJ. The content is solely the responsibility of the authors and does not necessarily represent the official views of the National Institutes of Health.

---

**Fig. S1.**
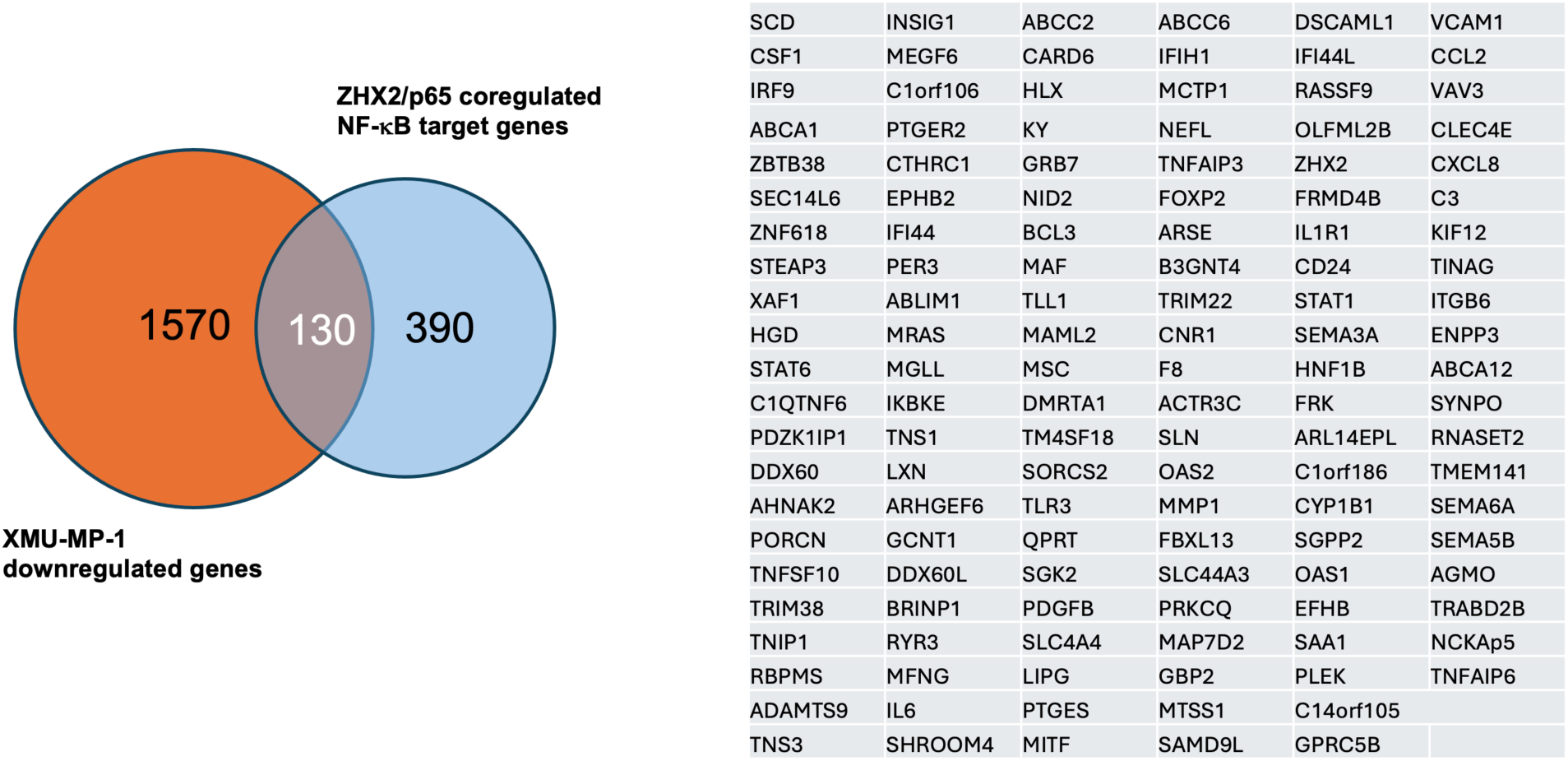
XMU-MP-1 inhibits NF-κB target genes co-regulated by p65 and ZHX2 Overlap of XMU-MP1 downregulated genes and NF-kB target genes co-regulated by ZHX2 and p65 in 786-O cells (left). List of the 130 NF-kB target genes co-regulated by ZHX2 and p65 and downregulated by XMU-MP1 in 786-O cells (right).

**Fig. S2.**
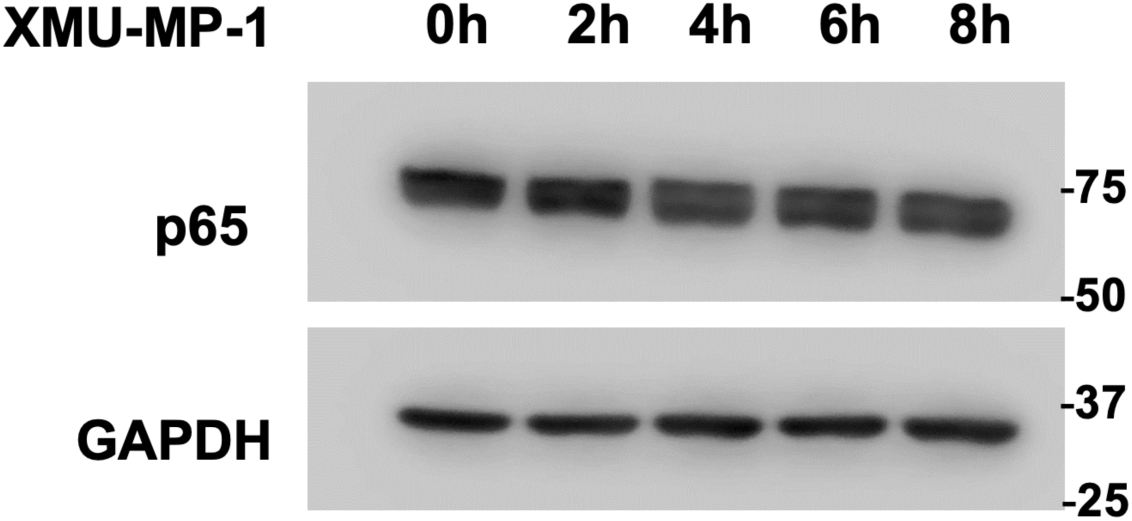
Hippo pathway inhibition does not affect p65 protein level Western blot analysis of p65 protein expression in 786-O cells treated with 2 μM XMU-MP-1 for the indicated time. GAPDH was used as a loading control.

**Fig. S3.**
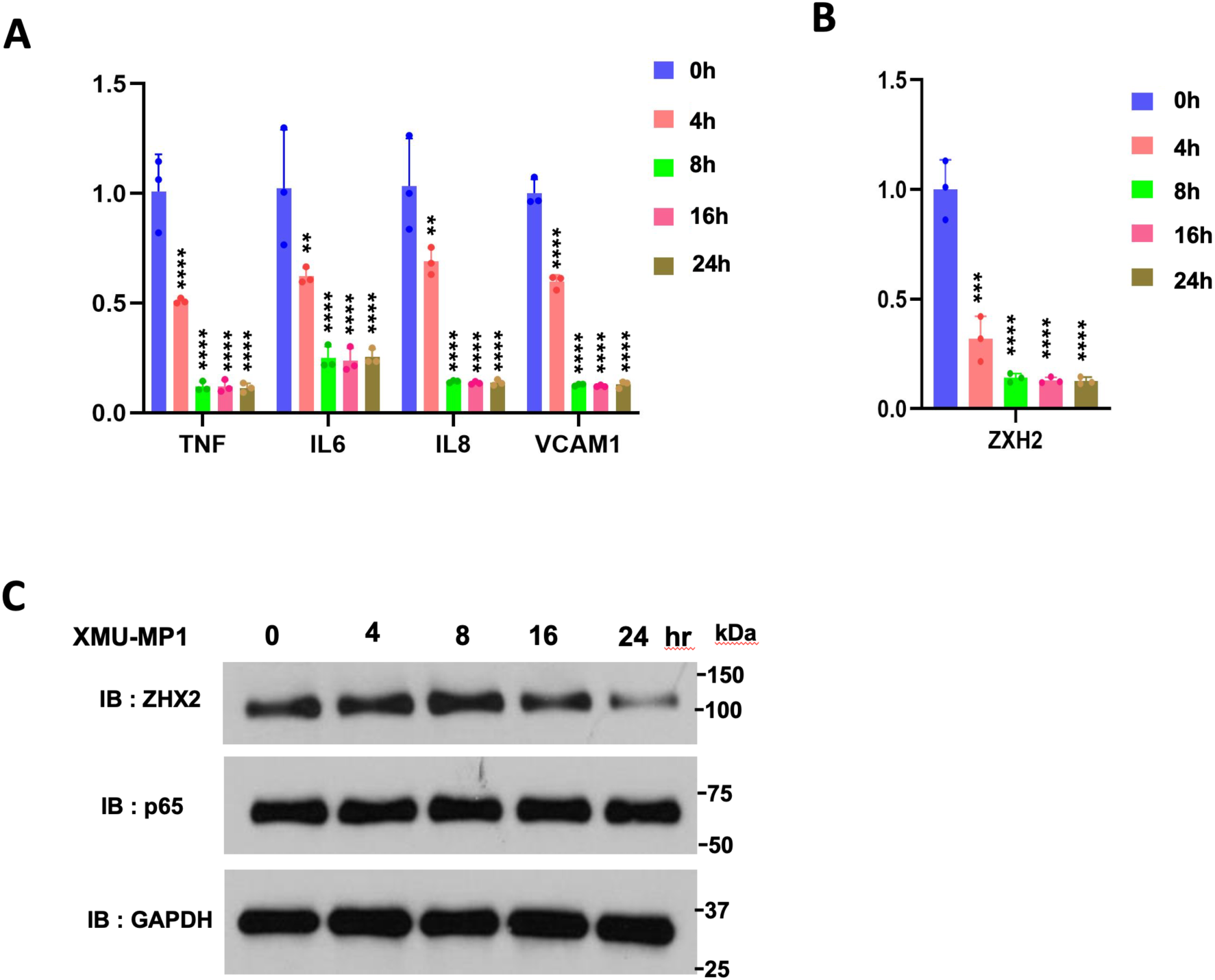
XMU-MP-1 inhibits NF-κB target gene expression without affecting ZHX2 protein level **A**-**B** Relative mRNA levels of the indicated NF-κB target genes (**A**) or *ZHX2* (**B**) in 786-O cells treated with 2 μM XMU-MP-1 for the indicated periods of time. Data in are ± SD. n=biological duplicates. **P<0.01, ***P<0.001, ****P<0.0001 (One-way ANOVA). **C** Western blot analysis of ZHX2 and p65 protein expression in 786-O cells treated with 2 μM XMU-MP-1 for the indicated periods of time. GAPDH was used as a loading control. ZHX2 level started to decline after XMU-MP-1 treatment for 16 hours while p65 protein level remained unchanged even after 24 hours’ treatment.

